# EPIGENOMIC VARIABILITY AND TRANSCRIPTOMICS AS A NOVEL MULTIOMIC COMPLEMENTARY APPROACH FOR PERSONALIZED NUTRITION IN COLORECTAL CANCER PATIENTS

**DOI:** 10.1101/2023.07.19.549686

**Authors:** Teresa Laguna, Oscar Piette-Gómez, Marco Garranzo, Marta Gómez de Cedrón, Ana Ramírez de Molina, Enrique Carrillo de Santa Pau

## Abstract

Food natural compounds are of interest as modulators of cancer progression and prognosis, as they participate in cellular processes such as growth and differentiation, DNA repair, programmed cell death and oxidative stress. Here we select dietary biocompounds for specific subgroups of 285 colorectal adenocarcinoma (COAD) samples by finding bioactives with opposite transcriptomic profiles to the subgroup-specific tumoral transcriptomes, hypothesizing they may counteract the cancer gene-expression profiles. To establish a CRC classification based on epigenetic variability, we selected 2,189 CpGs based on their differentially variable methylation between tumor and normal samples by a combination of linear and Bartlett tests. Samples were meta-clustered by 1) classifying each sample by 8 different methods (including k-means and hierarchical clustering), 2) building a network and 3) meta-clustering it by the *edge-betweenness* method. We extracted 6 main subgroups, 2 of them with immune-affected transcriptomes. We compared the transcriptomes of the 6 subgroups with the ones of 56 *in vitro* bioactive studies from GEO by Gene Set Enrichment Analysis (GSEA), resulting in a potential positive effect of resveratrol, japonicone A and vitamin D. In summary, we present a promising *in silico* strategy to suggest specific bioactives as co-adjuvants in cancer treatment.

## INTRODUCTION

Personalized nutrition is an area of growing interest, due to the known links between diet and human health (Balech et al. 2022; Livingstone et al. 2022; Ordovas et al. 2018; GBD 2019 Risk Factors Collaborators 2020) and specifically in human cancer (Sung et al. 2021). Several research groups have investigated the effect of modification of biocompound metabolism in the antitumoral drugs effectivity (Kanarek et al. 2018; Hopkins et al. 2018). However, little is known about the putative bioactive direct effect in the actual cancer cells or in the cell microenvironment, but it is believed that food compounds might only have minor to mild effects in cancer cell programmation (Shafabakhsh et al. 2019; Kanarek et al. 2020). However, it is also plausible that, due to tumor heterogeneity, those bioactive effects may be masked in some patients subgroups. Thus, designing specific bioactive supplementation for specific groups of cancer patients might enhance the drug therapy effects and reduce the effective dose which might reduce the secondary effects too.

Colorectal cancer (CRC) is the third most common cancer worldwide and second in death rate and has been extensively related to food and diet (Siegel et al. 2023; GBD 2019 Colorectal Cancer Collaborat.). There have been different efforts to classify colorectal patients in patient clusters to personalize treatments. The most popular classification is the CMS classification by the CRC subtyping consortium (CRCSC) (Guinney et al. 2015), by which samples were meta-clustered from 6 different classifications and research groups, based mainly on clinical and gene expression data. Although this classification is widely known, 13% of samples failed to classify in one of the 4 groups (CMS1-4). Characterization of these 4 subtypes were as follows: CMS1 was denoted as “the immune subtype”, with worse prognosis after relapse, high chromosome instability, mutations and methylator phenotype; CMS2 or the “canonical”, in which these samples show high copy number alterations; CMS3 was characterized by a metabolic deregulation but low copy-number and methylator phenotype, and CMS4 samples showed bad prognosis too and stromal infiltration and angiogenesis. In another study of the European consortia SYScol (Bramsen et al. 2017), different omics data were integrated by unsupervised class discovery through consensus non-negative matrix factorization (NMF)-based and samples clustered in 5 subtypes or “archetypes”: goblet, stroma, SSC (sessile serrated CRC), dARE (depleted in 3’ UTR AU-rich transcripts) and CIN (chromosomal instability), which accounted for clinical, transcriptomic and epigenomic data from epithelial and stromal tissue samples of >300 CRC patients. Although plenty of computational work was performed in these 2 studies, inter-sample epigenetic variation was not considered as a source of sample subtyping, not fully accounting for the possible differences in the tumor microenvironment (TME).

In this work we employ epigenetic variability as base for a new CRC classification, which has been previously described to be linked to cancer clinical diversity (Ecker et al. 2015; Hansen et al. 2011; Feinberg and Irizarry 2010), confirming previous findings that establish a correlation between epigenomic variability and CRC prognosis (Kasprzak and Adamek 2019; Molinari and Frattini 2013). Furthermore, molecular similarity analyses have been previously used to find disease comorbidities and drug repurposing (Sánchez-Valle et al. 2020; Fustero-Torre et al. 2021; Mateo et al. 2020). Here we incorporated those methodologies in a computational strategy with the aim to suggest personalized food supplements to specific CRC patient subgroups, which might be extended to other cancer types.

## MATERIALS & METHODS

### Data

Epigenomic data, sample-matching expression data and clinical & phenotypic data from Colon Adenocarcinoma patients (COAD) were obtained from the TCGA consortium. Specifically, level 3 - normalized DNA methylation data from 450k Illumina Methylation Arrays of 309 tumor samples and 37 normal adjacent tissues was downloaded from TCGA data portal (GDC). Matching transcriptomic data was obtained by downloading RNA-seq *fpkm* data from the same source. Samples not matching in their epigenomic and transcriptomic data were removed, resulting in 308 tumor and 19 normal adjacent tissues samples. Clinical data of common 327 samples included the variables *age, race/ethnicity, tissue of origin, ajcc stage* (AJCC Stage classification), *ajcc m, ajcc t, ajcc n* (AJCC TNM classification), *os* (overall survival), *pfi* (progression-free interval) and *dss* (disease-specific survival).

Transcriptomic data from 56 *in vitro* bioactive studies was obtained from GEO database (Edgar et al. 2002; Barrett et al. 2013; GEO - NCBI), which were selected based to include studies in transcriptomic arrays (Affymetrix, etc) with sample and gene annotations (table 1, supplementary file 1).

### Most variable CpG sites in COAD tumor samples

Differential variability of 450k-array probes between tumor and normal COAD samples was performed by the following methods: 1) *varFit* from *missMethyl* R package, which consists on fitting a linear model on Levene residuals of methylation M-values (Phipson et al. 2016; Phipson and Oshlack 2014), 2) *iEVORA* method (Teschendorff et al. 2016), based on simultaneously find differential methylation (t-student) and differential variability (Bartlett test). Common significantly variable CpG sites to both methods (127,853) were subjected to a bootstrap method to find the most variable CpGs. In each round, 37 random tumor samples were compared to the 37 normal samples and method *varFit* was applied. With n=500 repetitions, 28,188 CpGs were found as differentially variable in all rounds. To include the variability among tumor samples, absolute residuals of *M-values* of the bootstrap set were calculated and the 10% CpGs with most variable residuals were selected, resulting in a final set of 2,189 hypervariable CpGs sites. This 10% were empirically selected after observing variance distribution of the residuals (fig. S1).

### Patient / Sample Classification

Classification of the 308 COAD tumor samples was performed using the following different clustering methods based on the DNA methylation data of the selected hypervariable CpG set (2,189 sites). Specifically, an evaluation of best methods was carried out by means of the *clValid* R software package (Brock et al. 2008). This package employs both internal (Connectivity, Dunn, Silhouette metrics) and stability measures (average proportion of non-overlap (APN), average distance (AD), average distance between means (ADM), and figure of merit (FOM)) to evaluate the best clustering method and the optimal number of clusters. We compared hierarchical clustering, *k-means*, and CLARA methods, using 2-10 clusters, finding that the best methods were both the hierarchical clustering and *k-means* using 2 or 8 clusters.

To decide for a stable clustering, a meta-clustering based on the CMS classification (Guinney et al. 2015) was performed using the following 8 different clusterings:

- Beta-values 8 groups hierarchical
- Beta-values 8 groups k-means
- Beta-values 2 groups hierarchical
- Beta-values 2 groups k-means
- Absolute residuals 8 groups hierarchical
- Absolute residuals 8 groups k-means
- Absolute residuals 2 groups hierarchical
- Absolute residuals 2 groups k-means

Then, the Jaccard similarity among samples was calculated and a network built by using as distance: *dist = 1 - Jaccard* (fig S2) and removing the 60% of edges with the longest distances. A subsequent network clustering was performed using the *Fast-greedy* and *Edge-betweenness* methods of the *igraph* R software package, shown in fig S3. We selected a “core” group of samples of 285 forming clusters of >1 sample from the total of 308. Final clustering is shown in Figure 1, defining 6 clusters (DV2, DV3, DV4, DV6, DV7 and DV8). Final classification and phenotype of samples are shown in table 2 (supplementary file 1).

**Figure 1.**
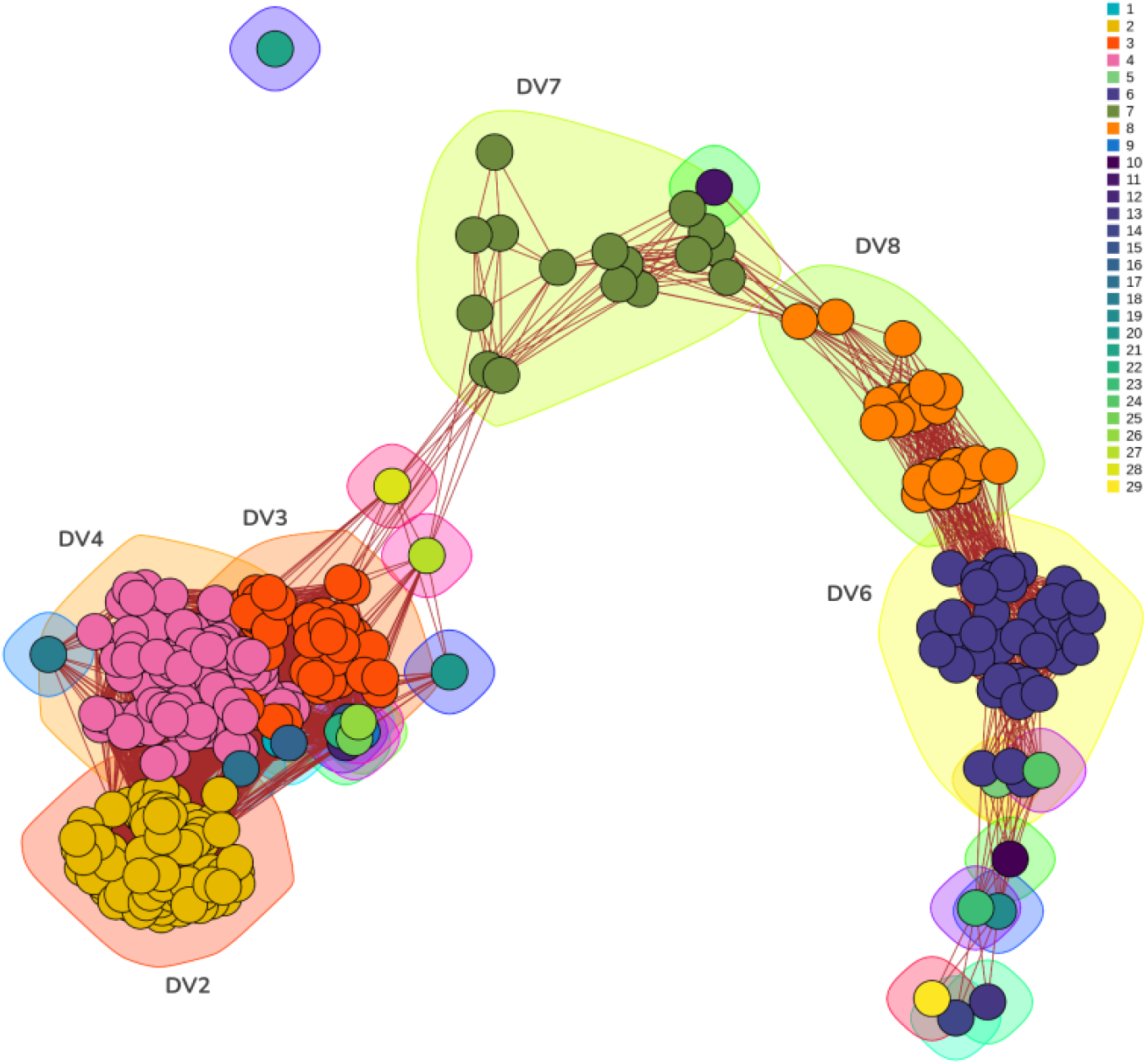
Colon Adenocarcinoma (COAD) subgroup classification based on epigenetic variability and network clustering. Network representing associations between COAD samples and sample clustering by the edge-betweenness method. Edges represent the closest 40% of distances between samples when *distance = 1 - Jaccard similarity* considering 8 different classifications per sample (see M&M).

### Phenotypic characterization of DV groups

We analyzed the multiple relationship between phenotypic variables, samples and categories by performing a Multiple Correspondence Analysis (MCA) by means of R packages *FactoMineR* and *factoextra*. For the enrichment analysis of sample categories in each DV subgroup, we performed individual hypergeometric tests per variable and per group. We established a probability for significance of over-representation to be >0.95 and for under-representation, to be <0.05.

### Analysis of differential expressed genes among DV groups

Differential expression analysis between each DV group and normal samples was performed using RNA-seq data from matching COAD samples with epigenomic data analyzed in previous sections, downloaded from the TCGA portal. Both *DEseq* and *limma* from respective R software packages were used (Love et al. 2014 and Ritchie et al. 2015, respectively) and common differentially expressed genes to both methods were selected (adjusted p-value by Bonferroni method < 0.05) (supplementary file 2).

### Pathway analysis

Pathway analysis of DV groups expression data was carried out by means of package *clusterProfiler* (Wu et al. 2021). Basically, the function *enrichGO* performs an hypergeometric test with the following options: *ontology = BP* (Biological Process), *pvalueCutoff = 0*.*05, qvalueCutoff = 0*.*10*. The gene universe comprised 30017 genes, and the ontology was obtained from ENSEMBL (org.Hs.eg.db).

### Personalized bioactive selection for each DV group

Transcriptomic data from arrays was analyzed by using the moderated t-test and empirical variance correction from the *limma* R software package (Ritchie et al. 2015). Top 250 up and down-regulated genes, based on the “*t*” statistic, were used as subsets to GSEA analysis (Subramanian et al. 2005), where absolute values of *log2(fold-change)* (*Log2FC*) of DV differential expression analysis were used to generate the gene rank for GSEA. “*t*” statistic was used to order the top 250 up and downregulated genes in bioactive studies because of: a) significance was almost null in all the 56 bioactive transcriptomics studies, b) *log2fc* ranges were very different among the 56 experiments, thus we considered “*t*” as a more stable metric. In the case of gene ranks of differential analyses of DV groups, we chose *log2FC* because all the groups belonged to the same experiment and metrics were consistent among the groups.

After performing GSEA, we considered significant analyses when NES >1.5 or NES<-1.5, which in all cases had a p-value <0.05, to standardize a similar value for all bioactive experiments.

## RESULTS

### New classification of Colon Adenocarcinoma patients based on epigenetic variability retrieved 6 main subgroups related with disease severity

Previous classifications of subgroups of Colon Adenocarcinoma are based mainly in gene expression data and clinical data (Guinney et al. 2015). We generated a new classification of COAD tumors based on variability of epigenomic data, as it has been related with cell lineages and tumoral activity (Suvà et al. 2013; Costa et al. 2023; Kim and Costello 2017). As tumor samples usually are a mix of different cell types and tissues, the presence of specific cell types can make tumors more aggressive or lethal (Hinshaw and Shevde 2019).

As specific DNA methylation levels in concrete CpG sites are characteristic of some cell types, to divide the TCGA COAD samples in subgroups, we investigated the most variable CpG sites: first, between tumoral and adjacent normal samples, and secondly among tumoral samples, selecting a subset of 2,819 CpGs (details in M&M). Optimal clustering of tumor samples was investigated, retrieving 8 different optimal classifications, which were used to calculate distance between samples by computing the difference to 1 of the Jaccard similarity (details in M&M). Distance among samples was used to build a network, removing the 60% longer edges. The sample network was clustered by 2 different methods: the fast-greedy (FG) method showed 3 main clusters (DV_FG1, DV_FG2, DV_FG3, Fig. S3) and the edge-betweenness (EB), 6 main clusters, defined as DV2, DV3, DV4, DV6, DV7 and DV8 (Fig. 1). FG classification contains EB classification, as the samples in DV2 are contained in the DV_FG1 subgroup, DV3 and DV4 samples in DV_FG2 and samples from DV6, DV7 and DV8 are classified as DV_FG3. A few samples were classified in individual clusters under EB clustering, but not using the FG method (fig.1 and fig. S3), thus we removed those of the EB classification and continued our analyses with the 285 core COAD samples.

### In-silico characterization of DV COAD subgroups

We retrieved phenotypical variables associated with each sample and patient from the TCGA databases, and evaluated their correspondence with the DV groups we have defined. For that, we performed a multiple component analysis (MCA) that showed a correlation of all three EB, FG and CMS classifications with the 2nd MCA dimension, and different categories of AJCC classification (Escrig Sos et al. 2019) with 1st dimension (fig. S4). Samples were also allocated in the 2-D dimensional space from MCA (1st and 2nd components, 15% of total variation - fig. S5). DV subgroups were distinguished in the first 2 components, being closer together the 3 DV groups DV6, DV7 and DV8, and the 3 groups DV2, DV3 and DV4 (fig. S6). Remarkably, the 1st component is associated most with Stage IV, metastasis state *M1* and disease-specific death (DSS) and the 2nd component with the cluster FG3 -*fast-greedy* classification-, DV6 (EB6) and CMS1 clusters, probably related with less severe disease (fig. S7). Whereas our classification based on DNA methylation variability does not show a prominent correlation with any AJCC-UICC classification (fig S8), we observed significant over- and under-representations of TNM categories in several DV subgroups (fig 2a-d). For example, TNM *M* classification is representative of DV subgroups, as DV2, DV4, DV7, DV6 and DV8 are enriched in samples in state M1a, M1, MX, M1b, and M0, respectively (fig.2a). In terms of tissue location of primary tumor, remarkably, DV2 and DV4 are enriched in tumors from the sigmoid colon, whereas DV3, DV6, DV7 and DV8 are enriched in tumors located in the cecum (fig. S9a).

**Figure 2.**
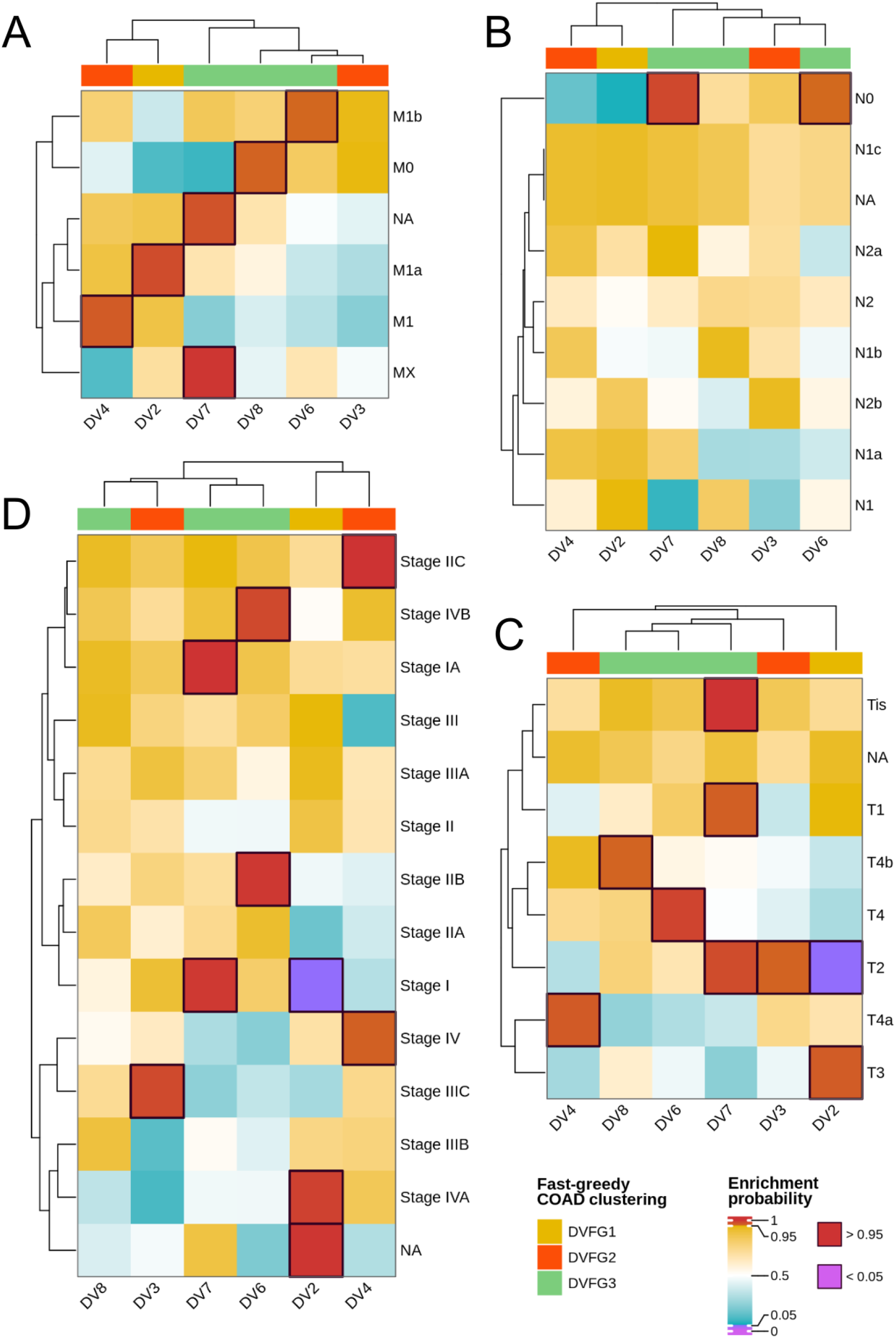
Enrichment of disease severity variables in DV subgroups. Heatmaps representing the results of hypergeometric tests among the proportions of each one of the 4 most relevant progression variables for colorectal cancer (CRC): (a) TNM classification “M”, (b) “N”, (c) “T” and (d) AJCC stage classification. Columns represent the DV subgroups; top annotation: the fast-greedy classification for each DV group; colors: probability of over-(>0.95 for significance) or under-(<0.05 for significance) representation.

Compared to CMS classification (Guinney et al. 2015), CMS2 and CMS4 samples were significantly enriched in DV2 and DV4 subgroups, whereas DV6 and DV8 were over-represented with CMS1 samples, and DV7 with CMS3 samples (Fig. S9b). No-labeled samples for CMS classification show enrichment in DV3 and DV8 subgroups (fig. S9b). A summary of phenotypic enrichment of each DV subgroup is shown in table 3 (supplementary file 1).

In general, DV2 and DV4 are enriched in samples with the most severe phenotypes, Stage IV, M1, T3/T4, with origin in the sigmoid colon and of CMS2 and CMS4 (fig. 2 and fig. S9). The only phenotypic differences are that DV2 is enriched in descendent colon samples and from younger people (<60 yr-old) and DV4 patients have received significantly more radiotherapy than the DV2 subjects. Although not significant, it is interesting to note that DV2 samples tend to be enriched in samples without progression and DV4 in samples which have progressed to worse outcomes (fig. S9h). In contrast, samples from DV7 are enriched in less severe phenotypes: Tis-T1-T2, Stage I, N0, originated in the cecum, and from alive and progression-free (and CMS3) patients. In DV8 are enriched samples with advanced stages but without metastasis. DV6 samples present a severe phenotype, but in this case with a specially dispersive outcome: T4, M1b, Stages IIb and IVb are enriched in this group, in addition to samples from CMS1, older people (>70 yr-old) and affecting ascending colon. DV3 subgroup constitutes an special group, enriched in samples from Stage IIIc and T2, and from the cecum like the less severe cases.

### Transcriptomic profiling of DV subgroups show specific functions enriched

We further characterized the 6 DV groups by analyzing differences in transcriptomes from the 308 COAD methylation-matched RNAseq samples. We found 3,683 genes differentially transcribed between any DV subgroup and normal samples and used their RNA-seq signal to compare clustering classifications. As expected, there was no significant correlation between the 3,683-RNAseq signal to the DV classification, and barely a limited correlation with CMS classification (fig. S10).

Nonetheless, we observed several differences when we investigated for enriched-pathways in transcription of each group compared to normal samples. We used the differential expression analysis data from the 308 samples to perform an enrichment pathway analysis based on the Gene Ontology database (Gene Ontology Consortium et al. 2023). Pathway analysis showed a general up-representation of cancer-related pathways in all groups, such as muscle system, blood circulation and ion transport pathways (fig. 3, fig. S11, fig. S12 and supplementary fig. 3). Looking at specific routes, immune function-related pathways were overrepresented in DV4 and DV6 groups, with a significant downregulation of genes involved in immunoglobulin production (fig. 3, fig. S11 and supplementary fig. 3). Furthermore, DV2, DV3 and DV4 transcriptomes are enriched in hormone metabolism and extracellular matrix organization processes, whereas DV7 and DV8 show enrichment in synaptic-related pathways (fig. 3, fig. S11 and supplementary fig. 3). These results suggest immune ablation in specific DV subgroups, although not related to tumor severity.

**Figure 3.**
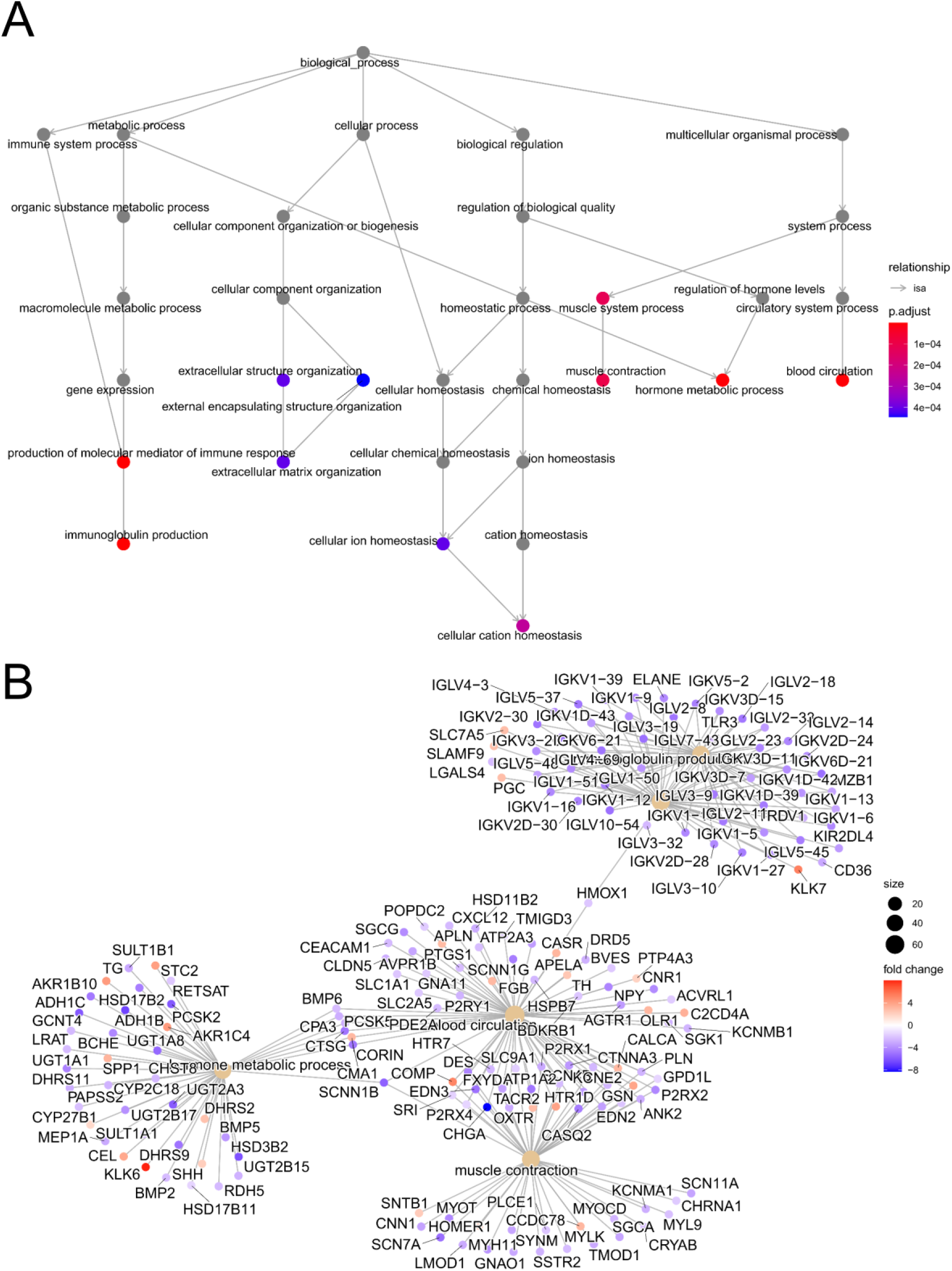
Biological pathway overrepresentation in gene expression of DV4 group compared with normal samples. a) Over-represented routes displayed in a parent-child tree. Significant over-represented pathways are displayed in color (see legend). b) Over-(red) and under-(blue) expressed genes and the main pathway which they belong to displayed in a network graph.

### Resveratrol, japonicone A and withaferin A as potential beneficial bioactives with general effect and vitamin D with specific action in “immune” COAD subgroups

After CRC DV subgroup characterization, we explored the concept of transcriptional-based drug repurposing (Jahchan et al. 2013; Iorio et al. 2010; Sánchez-Valle et al. 2020; Mateo et al. 2020), to suggest bioactive compounds as potential directed supplementations in specific COAD DV-patient subgroups. We investigated bioactives with opposite transcriptomic effect to the cancer transcriptomic signatures of each patient group considering the hypothesis that they might ameliorate the cancer expression profile or enhance the anti-tumor program (Mateo et al. 2020). We performed Gene Set Enrichment Analysis (GSEA) (Subramanian et al. 2005) in each subgroup gene expression profile considering as gene subsets the top 250 up- and down-regulated genes from 56 transcriptomic bioactive studies. Considering p-value<=0.05 for significance of the GSEA analysis, we obtained 46 bioactive conditions with a potential beneficial effect at the transcriptomic level to at least 1 DV group (Fig. 4, table 4 supplementary file 1).

**Figure 4.**
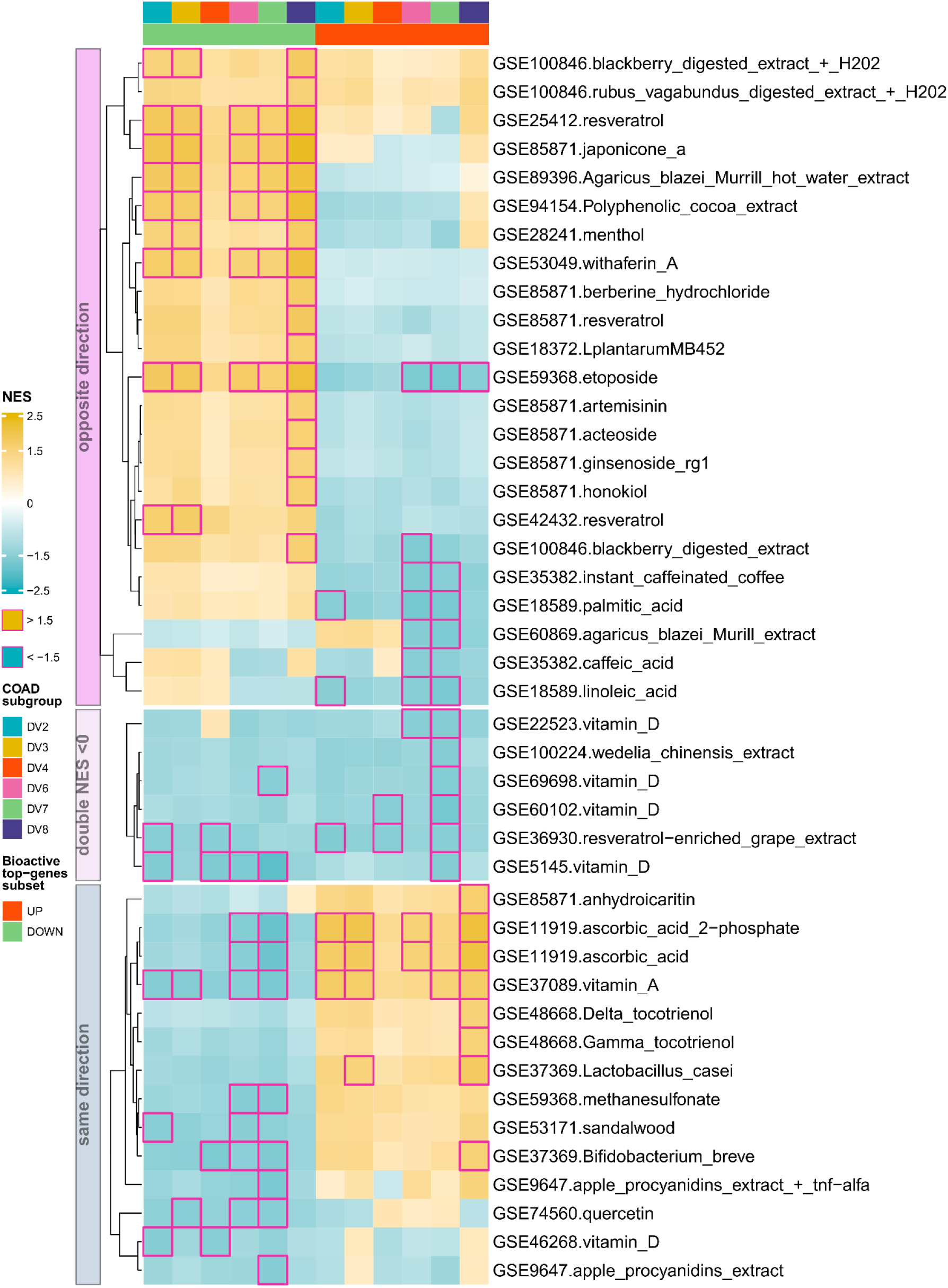
Proposed bioactive compounds to have a positive or negative effect in at least one of the 6 COAD subgroups. Heatmap representation of Gene Set Enrichment Analysis (GSEA) of top 150 over-(columns, red) and under-expressed (columns, green) genes of a bioactive study (rows) in each Dv subgroup transcriptomic profile (columns). Significant enrichment was considered when NES >1.5 or NES > -1.5 (denoted by a square in the corresponding cells).

Out of the 46, resveratrol, japonicone A, *Agaricus blazei* Murrill extract, polyphenolic cocoa extract and withaferin A are potential candidates to counteract all DV groups (except for DV4) *cancer expression signature* by hypothetically down-regulating highly expressed genes, as they show a significant Normalized Enrichment Score (NES) >0 of their most down-regulated genes. For palmitic and linoleic acids, their up-regulated subset shows a significant NES<0 with DV2, DV6 and DV7 transcriptomic profiles. Remarkably, several vitamin D studies show negative NES in their up-regulated subset -making it a candidate for design of its supplementation- and/or in their down-regulated genes (Fig. 4, “double NES<0”), therefore, their potential beneficial effect might be limited. Its putative counteracting effect might be present in DV4, DV6 and DV7, suggesting an activation of the immune functions. Interestingly, ascorbic acid (vitamin C) and vitamin A might elicit a transcriptomic profile with similar tendencies to the DV transcriptomes, as their up-regulated subsets show a significant NES>0 and their down-regulated subsets, a NES >0 (Fig. 4, “same direction”).

## DISCUSSION

Here we have presented a general *in silico* strategy to generate lists of potentially beneficial biocompounds to specific CRC cancer subgroups, classified according to epigenetic variability.

Several efforts have been made to define colon cancer subtypes based on integration of omics data, such as the popular CMS classification (Guinney et al. 2015) and the SYScol classification (Bramsen et al. 2017). However, these classifications rely on the bulk-nature of samples, which may vary enormously in cell type composition and tumor purity (Aran et al. 2015; Zheng et al. 2017). CMS classification considers gene expression as the main driver of subtype classification, which may be affected by a plethora of factors. In contrast, epigenomic landscape tends to be more stable and characteristic of cell type, including of tumoral types, which show an increased variability in DNA methylation. Here, interestingly we have defined a new subgroup classification of the colon cancer patients based on DNA methylation variability which reflects phenotype and severity diversity, so personalized treatments and food supplementation might be directed to each one of these groups. Through our network clustering of samples, we may infer that from the samples with less severe outcomes, which characterize DV7 subgroup, cancer progression may evolve in 2 directions: one to the DV8-DV6 subgroups, and the other to the DV3-DV2-DV4 subgroups. Further molecular and experimental characterization would be required to confirm these 2 progression ways. DV4 and DV6 samples, the ones with the worst prognosis, show an expected down-regulation in immune pathways, especially in the immunoglobulin production. DV2 and DV8 represent samples with intermediate-severe prognosis but without metastasis in the case of DV8. DV3 subgroup presents an intermediate progression state, which may be more prone to be helped, in addition to DV7 samples, with food supplementation.

Recently, several pieces of evidence show the interference of diet compounds in anti-tumoral therapy (Kanarek et al. 2018; Collado-Borrell et al. 2016). In addition, effective immune function is generally impaired in cancer due to immune evasion (Jhunjhunwala et al. 2021; Tauriello et al. 2018). Both factors for tumor growth and expansion might be counteracted by the supplementation of specific biocompounds in the patients diet. From the bioactives with potential beneficial effect to cancer expression onset, i.e., the bioactives with NES >0 in their down-regulated subset and/or NES<0 in their up-regulated subset, we can divide them in 2 groups. First, bioactives with a potential “inhibitory” effect (NES>0 in their down-regulated genes) in all subgroups (except in DV4): here there are japonicone A, resveratrol, polyphenolic cocoa extract and withaferin A. On the other side, there are bioactive with a putative “stimulatory” effect (NES<0 in their up-regulated genes) but potentially benefit the DV7-DV8-DV6 branch: palmitic and linoleic acids, coffee-related bioactives, blackberry digested extract. To support the potential of the GSEA method to personalized nutrition, we found an important significant inhibitory and stimulatory putative effects of etoposide, which is a chemotherapeutic essayed in one of the 56 bioactive studies and included here to show the potential of the analysis. Several bioactives have already been described to show anti-tumoral effects, such as japonicone A, resveratrol, and vitamin D. The role of vitamin D as stimulatory of immune functions may be limited by the inhibitory effect of other genes, as it shows a bivalent behavior in GSEA. However, it is the only bioactive which we found a potential positive action in DV4, in addition to DV6, the subgroups with the worst prognosis.

Here we present a computational workflow to suggest bioactive compounds and/or foods for specific subgroups of colorectal cancer patients, with the aim to improve the patient quality of life and disease evolution. This strategy might be extended to other cancer types, always considering that the potential positive effects of the suggested bioactives has to be experimentally validated before a nutritional intervention study. Vitamin D and the rest of the bioactives proposed here are dependent on the bioavailability of their molecular integrity in the target cells, which may be the tumoral or not, as the bioactives assays were performed in different cell lines from not only colon tissues. The counteracting molecular effect is also an hypothesis that requires further experimental validation. Nonetheless, with the growing availability of molecular data of bioactive effects, we consider this as the first stone to automatize personalized nutrition for a broad spectrum of cancer patients.

## ACKNOWLEDGEMENTS

The results shown here are in part based upon data generated by the TCGA Research Network: https://www.cancer.gov/tcga. We would like to thank Dr. Andrew Teschendorff and team for the iEVORA method and also Vera Pancaldi and Simone Ecker for their helpful discussions. Supported by Spanish Plan Nacional I+D+i PID2019-110183RB-C21 and partially by Food Nutrition Security Cloud (FNS-Cloud), which has received funding from the European Union’s Horizon 2020 Research and Innovation programme (H2020-EU.3.2.2.3. – A sustainable and competitive agri-food industry) under Grant Agreement No. 863059.

