## Supplementary Figures for "EPIGENOMIC VARIABILITY AND TRANSCRIPTOMICS AS A NOVEL MULTIOMIC COMPLEMENTARY APPROACH FOR PERSONALIZED NUTRITION IN COLORECTAL CANCER PATIENTS"

Laguna et al.

# S1

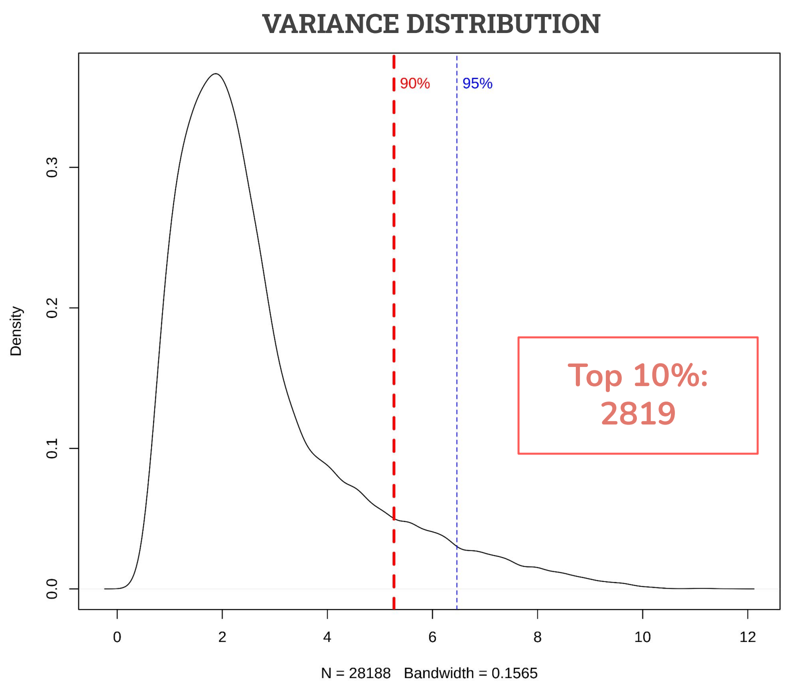


**Supplementary Figure 1. M-values variance distribution from the 28,189 top variable CpG sites**. We had previously calculated the most variable CpG sites between the 305 TCGA COAD tumor samples and their 37 normal samples counterparts. To select the most relevant subset we plotted the residuals distribution only in tumor samples. The 10% values with highest variance were selected for 2 reasons; 1) 2,819 CpGs sites is a manageable number to work with for a signature, and 2) we observed the elbow in the distribution and empirically decided the top 10% does not correspond to the null distribution.

# S2

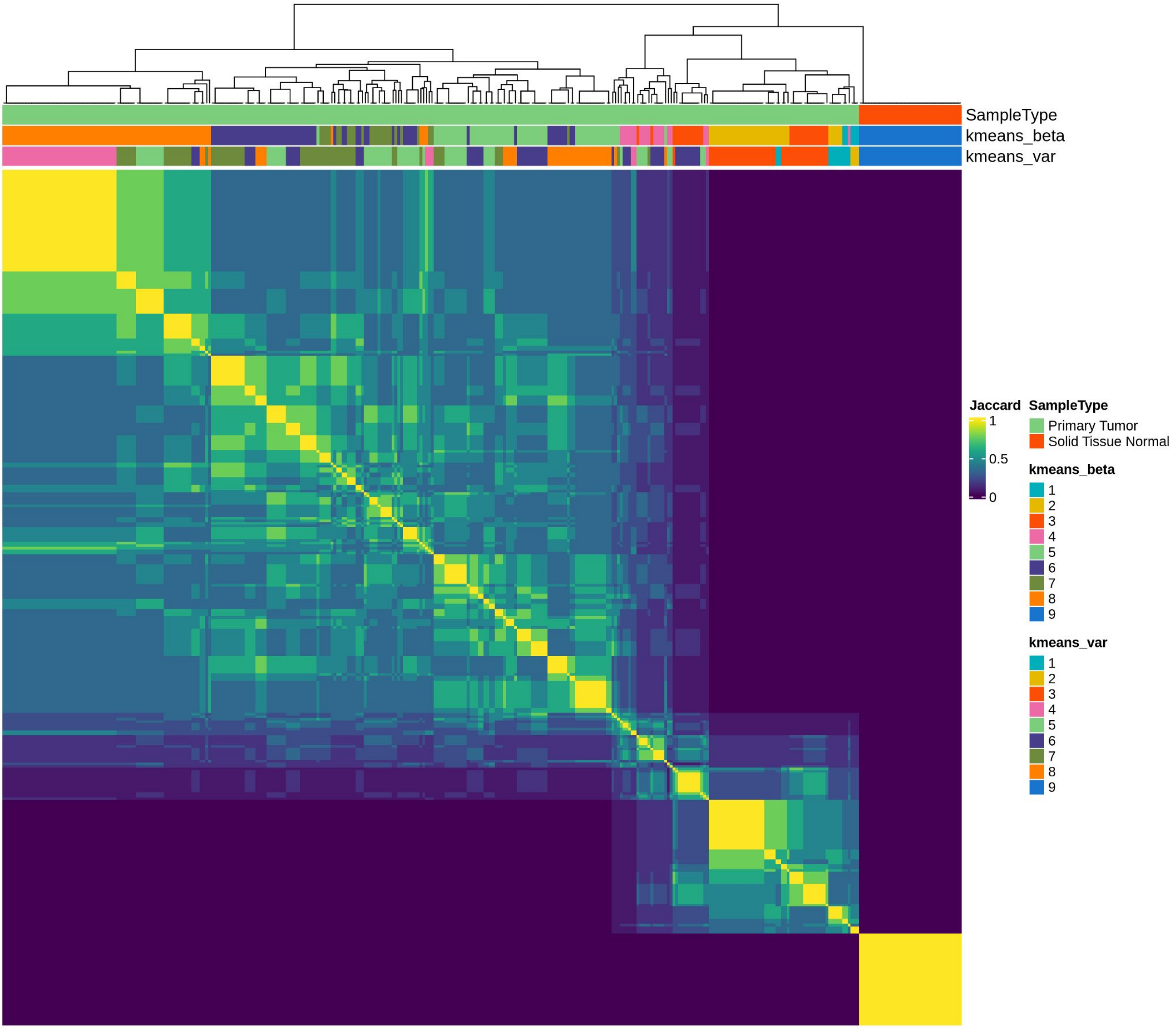


**Supplementary Figure 2. Jaccard similarity between the 343 COAD samples (308 tumor and 37 normal)**. Each one of the 342 samples was classified according 8 different clustering methods, all of them based on the methylation values of the selected 2,819 most variable CpGs. 2 of these classifications were generated by *k-means* using the beta values or the residuals (top annotation). Jaccard similarity was calculated for each pair of samples and represented in a correlation plot, in which samples are ordered similarly in rows and columns.

# S3

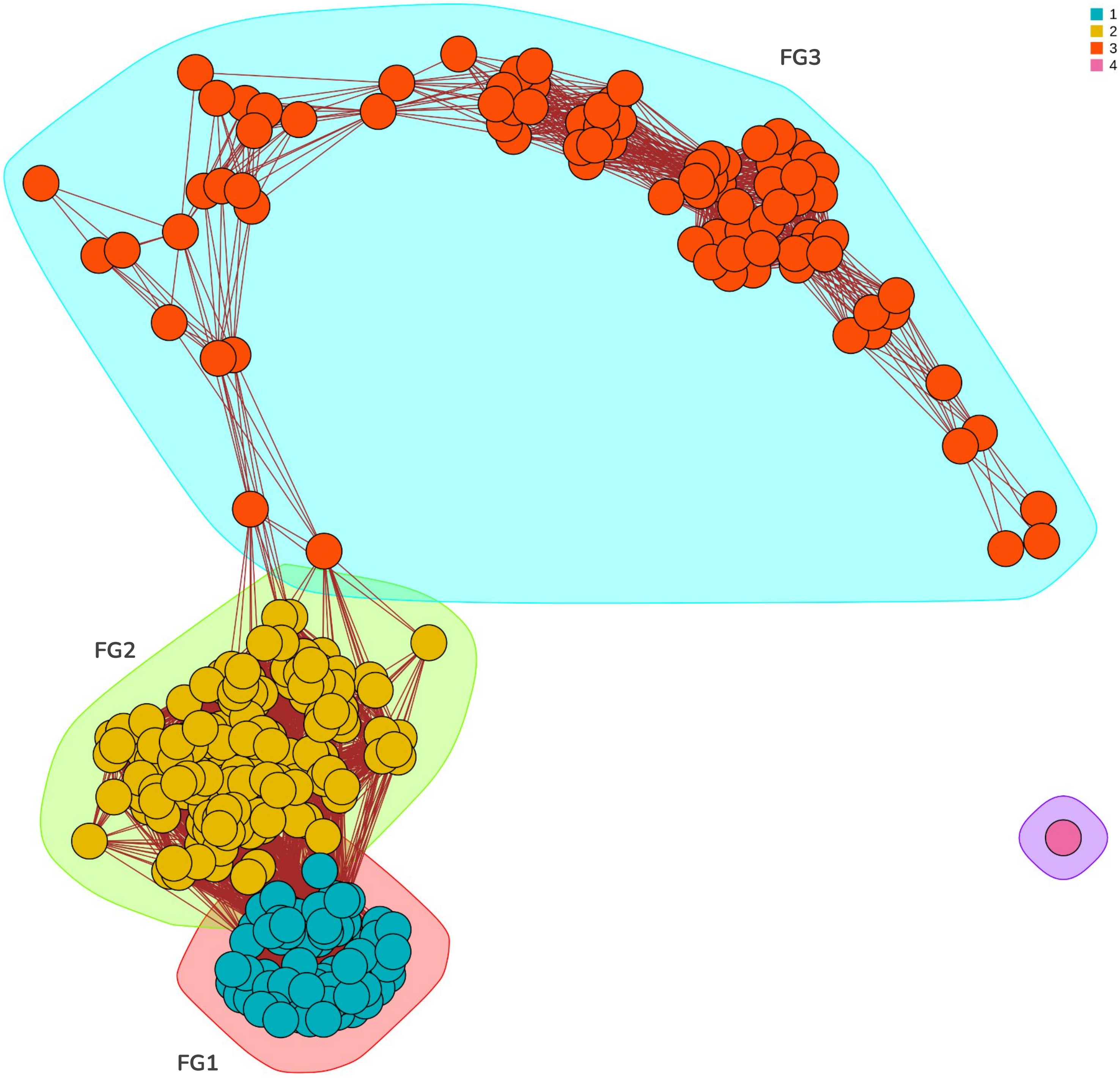


**Supplementary Figure 3. Network-based meta-clustering of the COAD samples by the *fast-greedy* method.** Distance between samples was calculated by the formula *1 - Jaccard similarity* and 60% of longer distances were removed. 3 major clusters were obtained, plus an individual cluster,

# S4

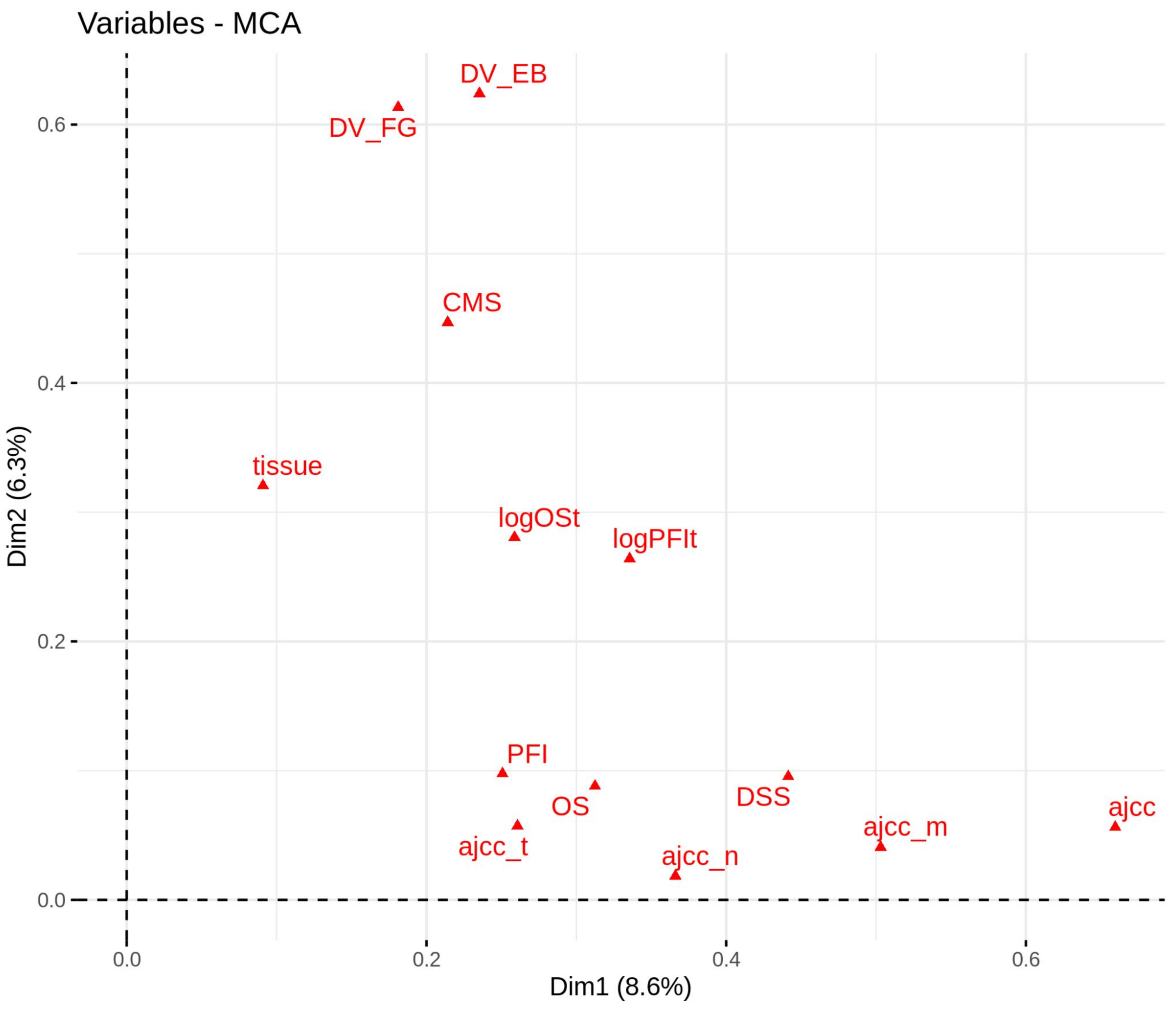


**Supplementary Figure 4. Correlation of phenotypic variables with the first 2 components of the MCA analysis.** Multiple correspondence analysis was performed for phenotypic variables and the 2 meta-clustering classifications developed in this manuscript. *DV_EB*: meta-clustering by the *edge-betweenness* method; *DV_FG*: meta-clustering by the *fast-greedy* method; *CMS*: CMS classification; tissue: tissue of tumor origin; *logOSt*: ln of Overall Survival time; *logPFIt*: ln of Progression Free Interval time; *PFI*: Progression Free Interval (bimodal 0/1); *OS*: Overall Survival; *DSS*: Disease Specific Survival; *ajcc_t*: “T” category of AJCC TNM classification; *ajcc_n*: “N” category of AJCC TNM classification; *ajcc_m*: “M” category of AJCC TNM classification; *ajcc*: AJCC Stage classification.

# S5

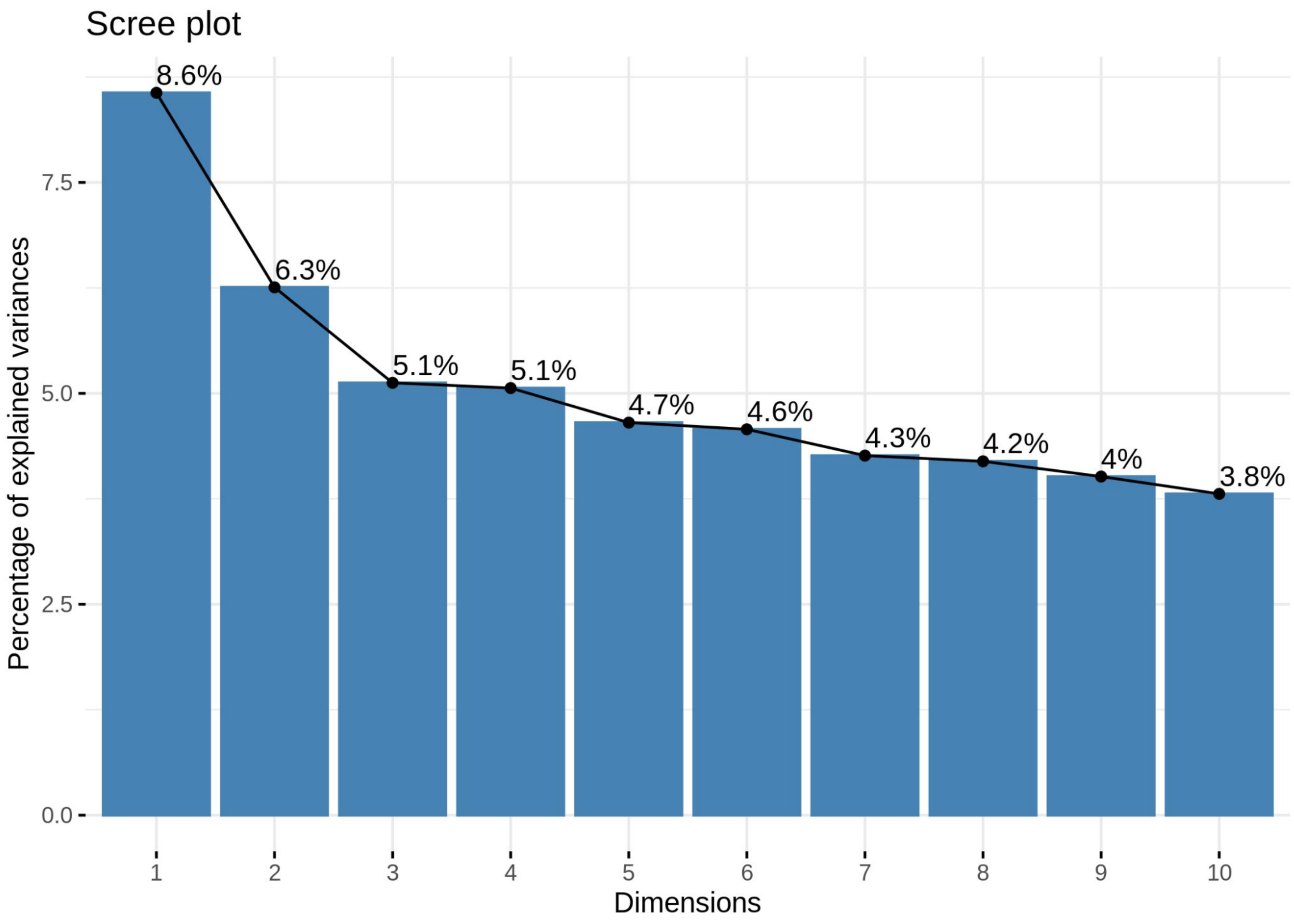


**Supplementary Figure 5. MCA analysis scree plot.** Barplot representing the percentages of variances explained for each one of the first 10 components/dimensions of the MCA.

# S6

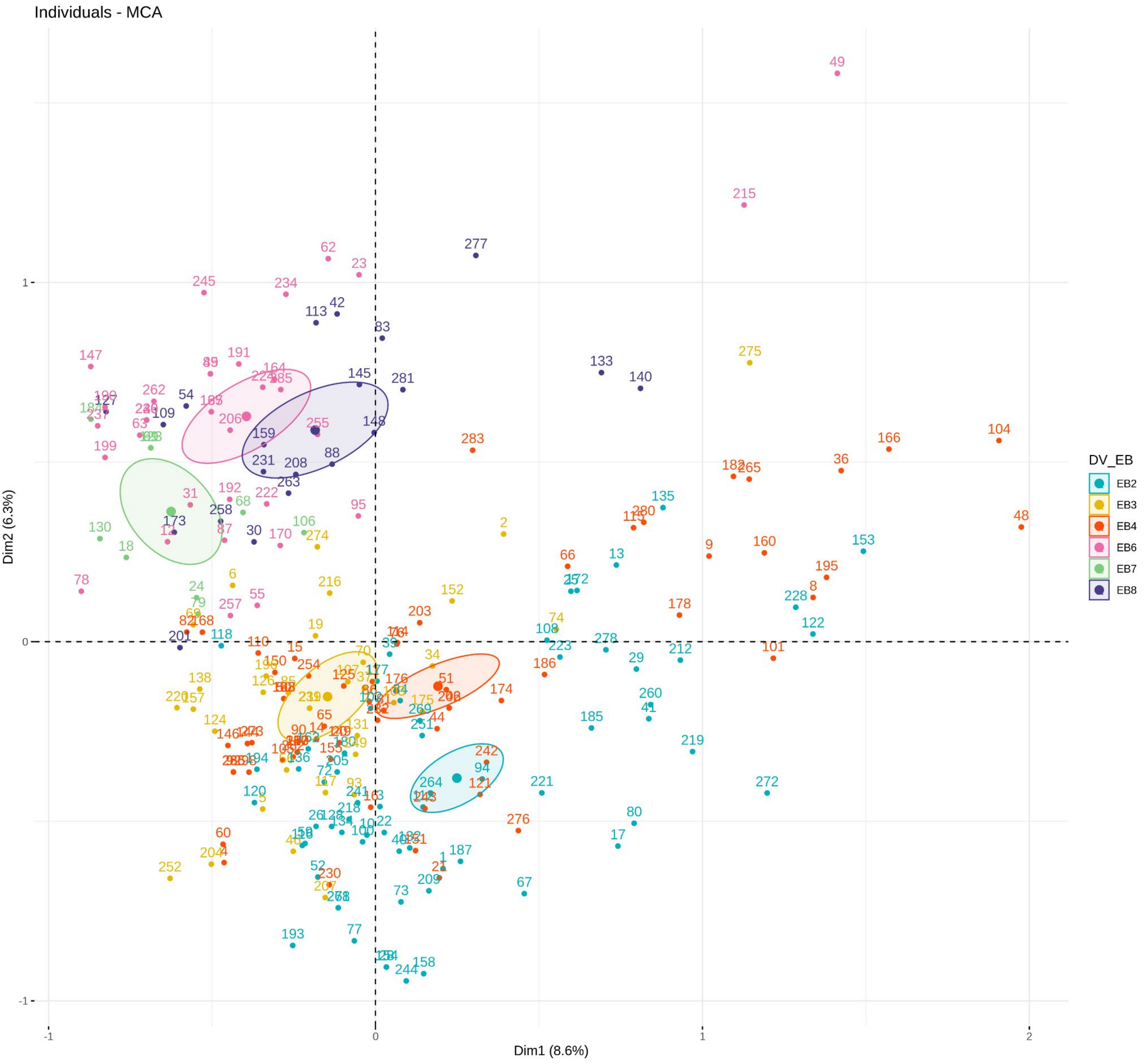


**Supplementary Figure 6. Sample coordinates in the first 2 components of de MCA analysis.** Samples are colored by their DV_EB (differential variability edge-betweenness) clustering. Same coloring applies for cluster centers and 95% confidence ellipses.

S7

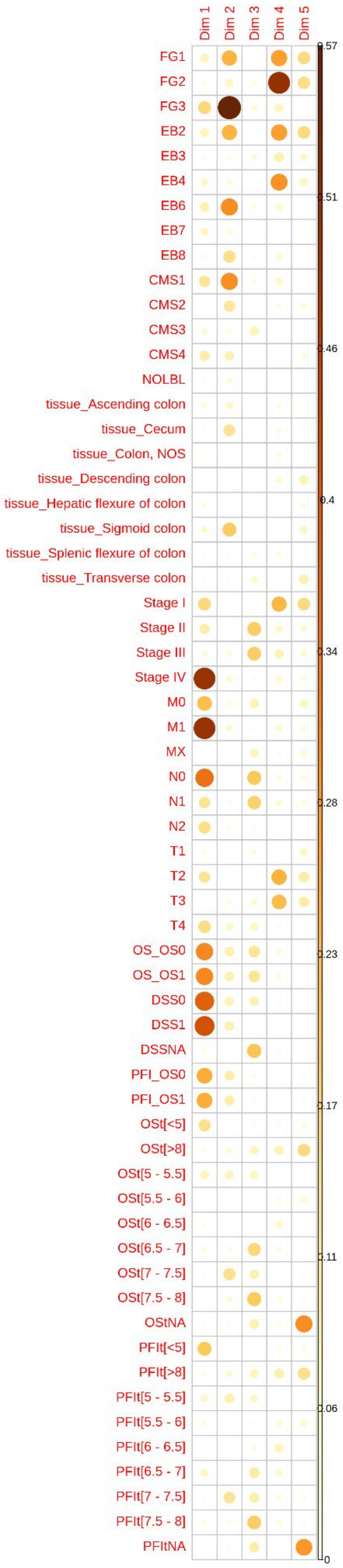


**Supplementary Figure 7. Correlation between variable categories and the first 5 components of the MCA analysis.** Color and diameter of spheres refer to the cos2 value of each pair category-component.

# S8

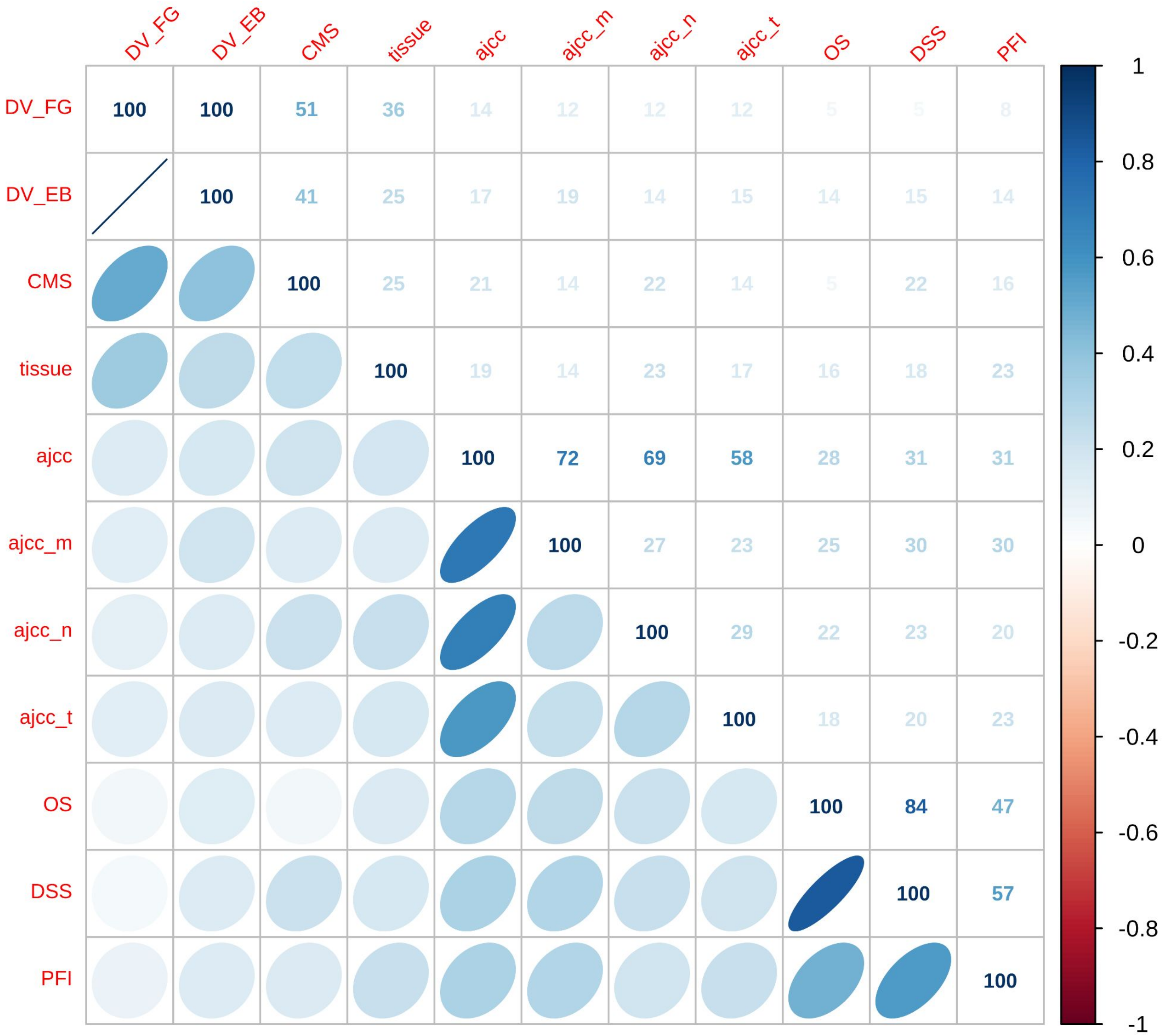


**Supplementary Figure 8. Correlation between pairs of variables from the MCA analysis.** Values of correlation are denoted by a numeric percentage (over the diagonal) and a distribution ellipse (under the diagonal). Color indicates the intensity of the correlation. *DV_EB*: meta-clustering by the *edge-betweenness* method; *DV_FG*: meta-clustering by the *fast-greedy* method; *CMS*: CMS classification; tissue: tissue of tumor origin; *logOSt*: ln of Overall Survival time; *logPFIt*: ln of Progression Free Interval time; *PFI*: Progression Free Interval (bimodal 0/1); *OS*: Overall Survival; *DSS*: Disease Specific Survival; *ajcc_t*: “T” category of AJCC TNM classification; *ajcc_n*: “N” category of AJCC TNM classification; *ajcc_m*: “M” category of AJCC TNM classification; *ajcc*: AJCC Stage classification

S9

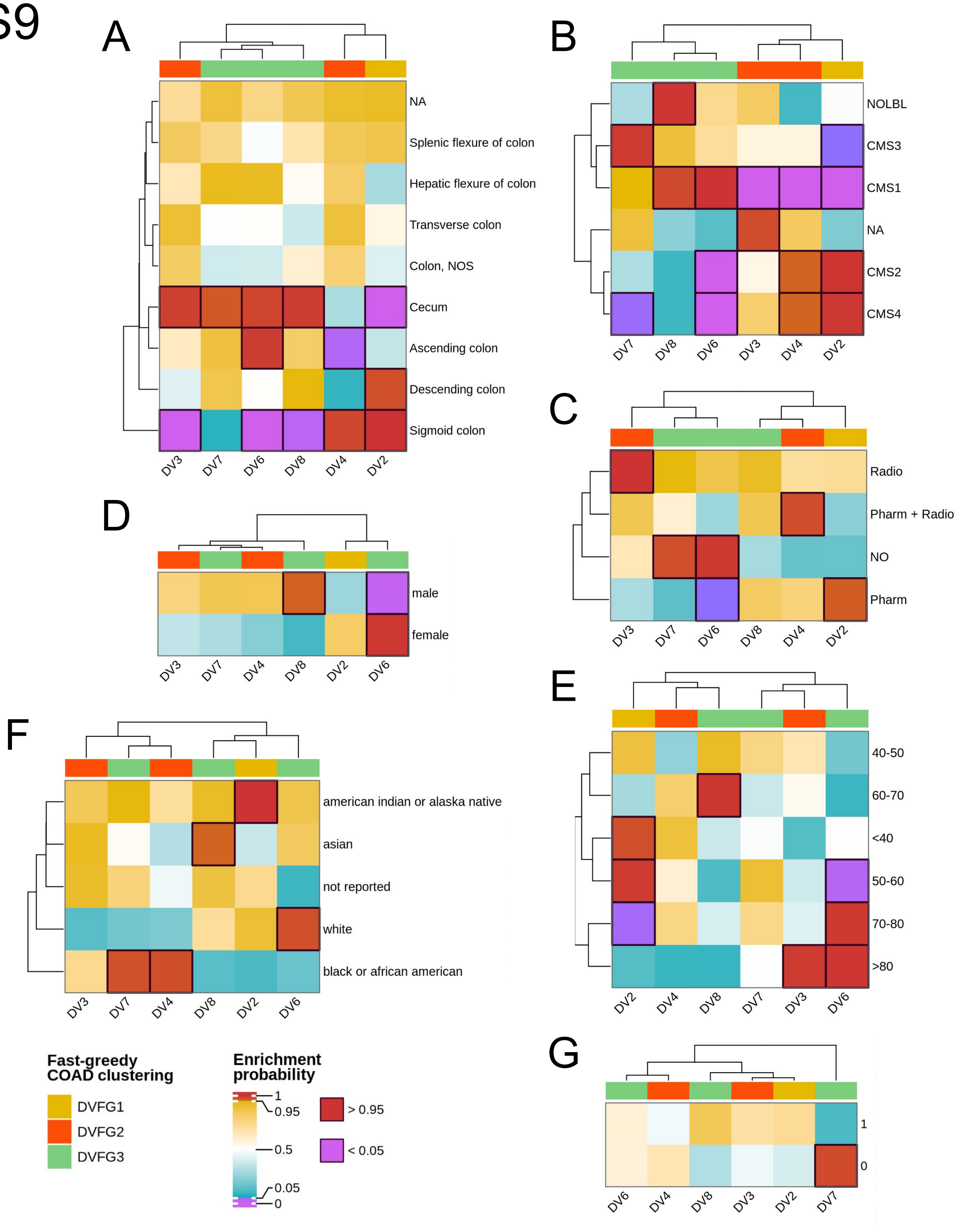


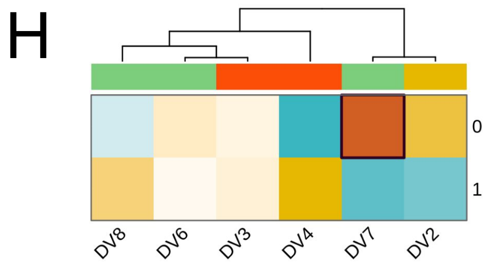


**Supplementary Figure 9. Enrichment of phenotypic variables in DV subgroups.** Heatmaps represent the results of hypergeometric tests among the proportions of: (a) tissue of origin, (b) CMS classification, (c) treatment, (d) sex, (e) age group, (f) race/ethnicity, (g) OS, (h) PFI. Columns represent the DV subgroups; top annotation: the fast-greedy classification for each DV group; colors: probability of over- (>0.95 for significance) or under- (<0.05 for significance) representation. Cells with significance are squared.

# S10

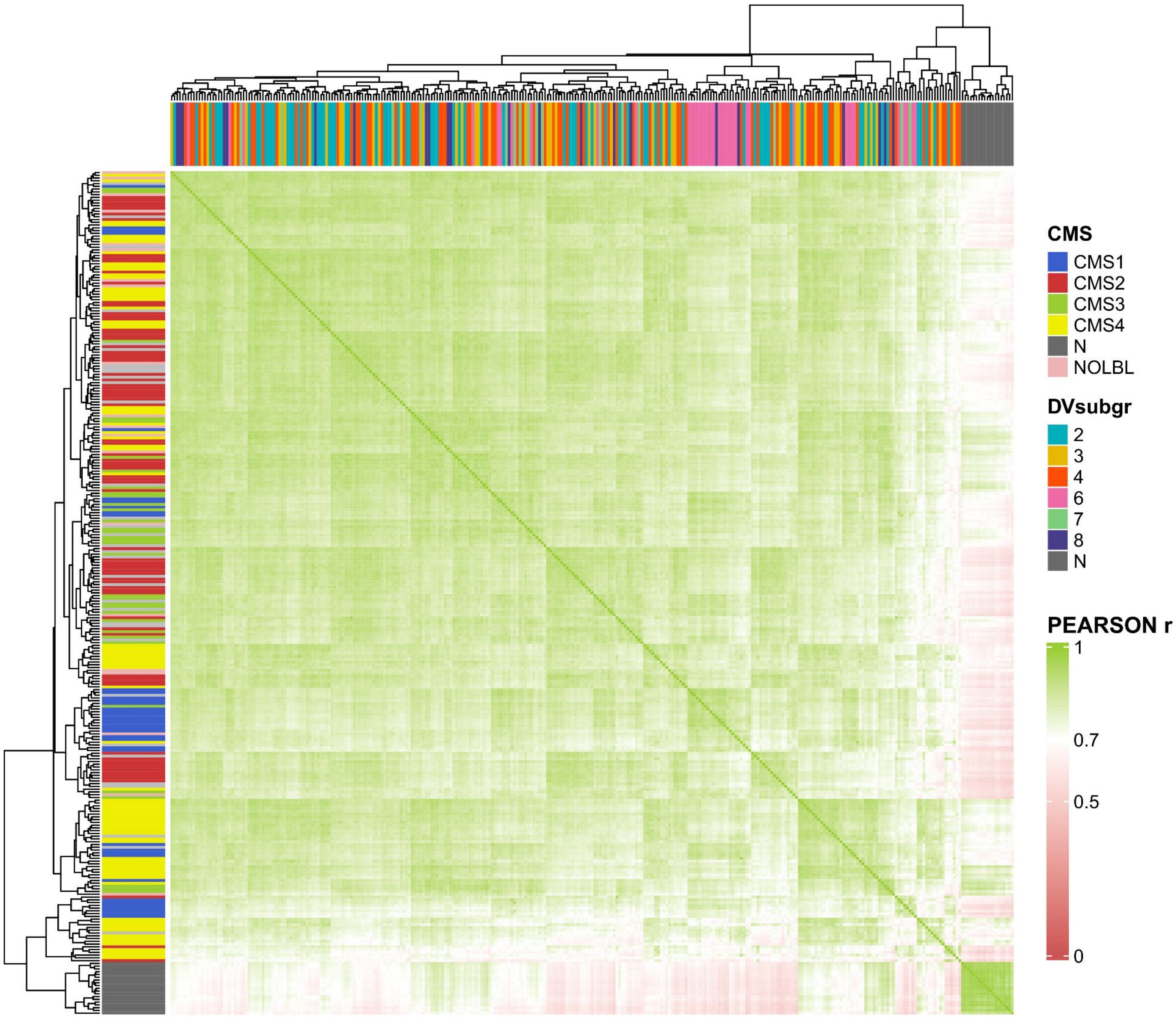


**Supplementary Figure 10. Correlation between pairs of COAD samples based on differentially expressed genes.** Normalized expression of the 3,683 differentially expressed genes between any DV group and normal samples was employed to plot the *Pearson r* correlation coefficients. Top annotation, sample category in DV_EB classification; Left annotation: sample category in the CMS classification.

# S11

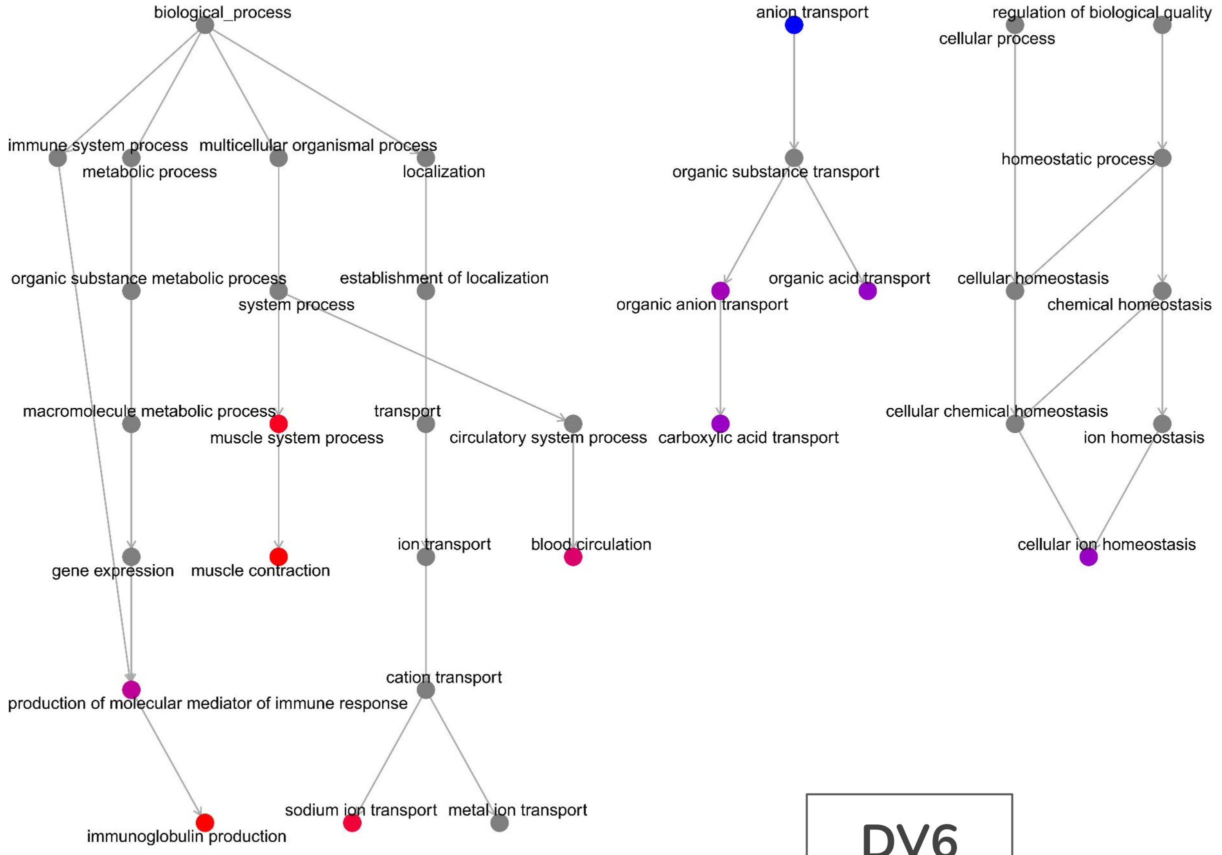


DV6

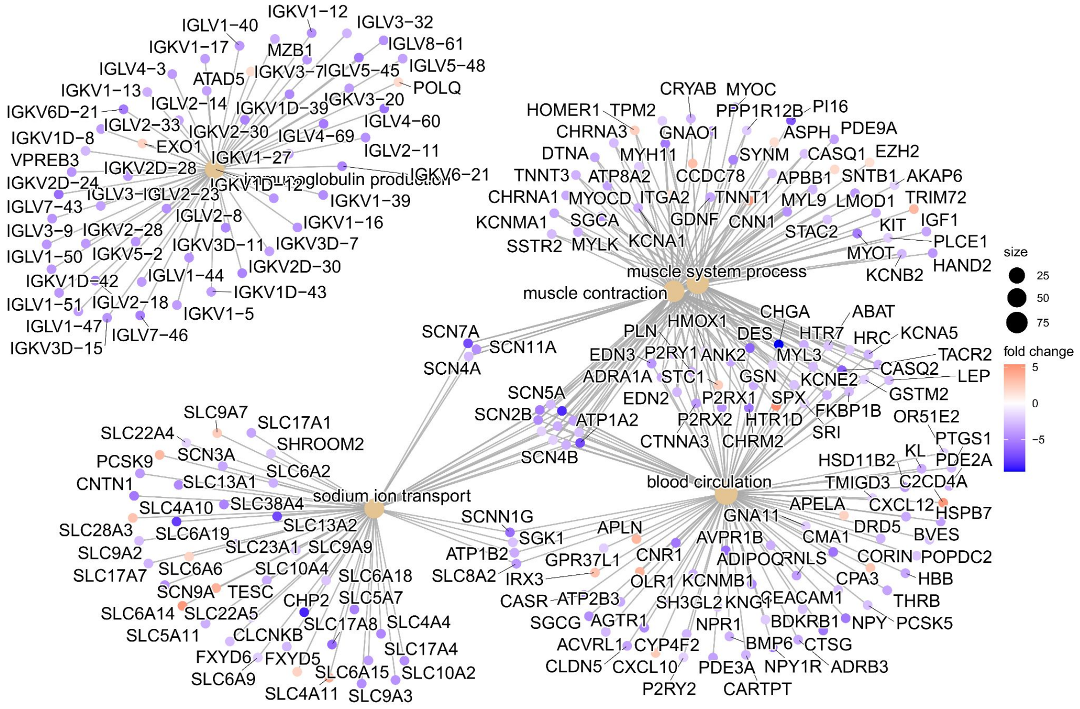


**Supplementary Figure 11. Pathway over-representation analysis for samples in DV6 group.** Significant results of hypergeometric tests using as subsets differentially expressed genes between DV6 and normal groups and as ontology *Biological Process* category from Gene Ontology Consortium. *Top*, significant routes (colored) displayed in a parent-child tree configuration. *Bottom*, differentially expressed genes members of the most significant over-represented pathways, displayed in a network configuration.

# S12

## DV2

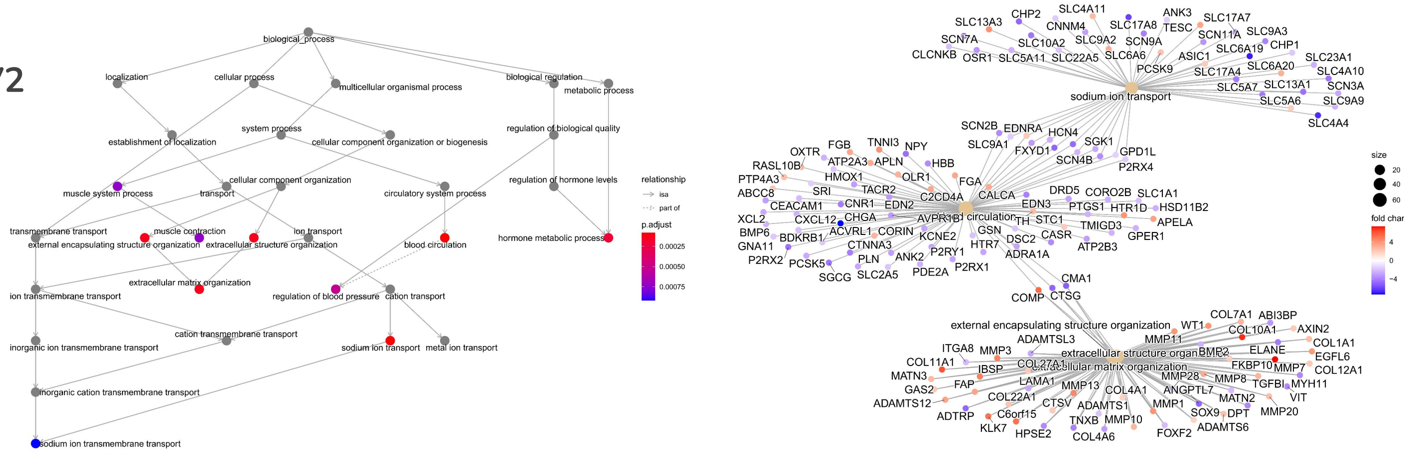


## DV3

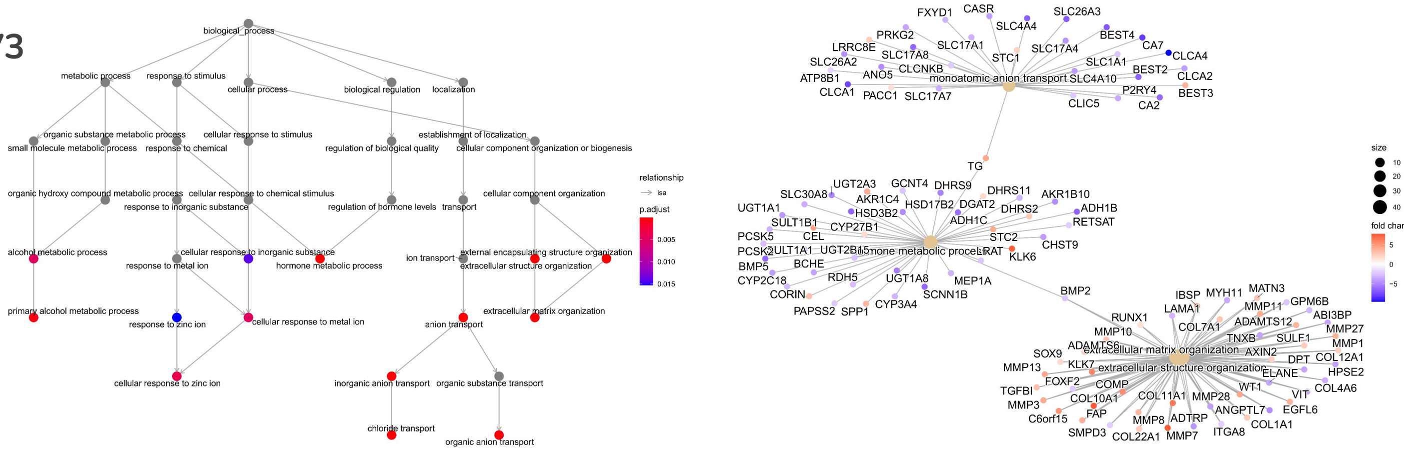


## DV7

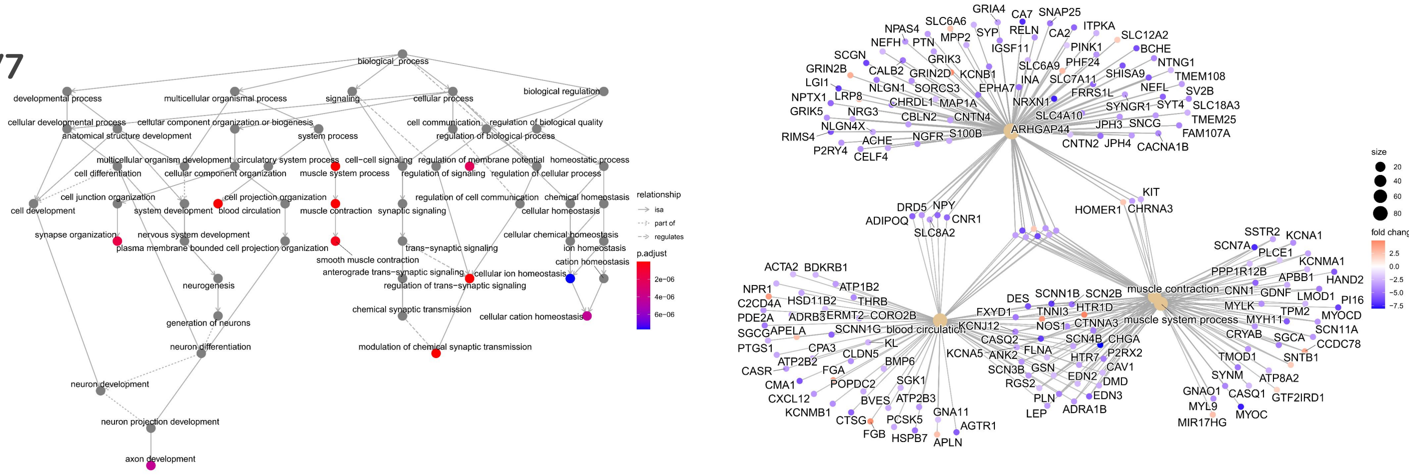


## DV8

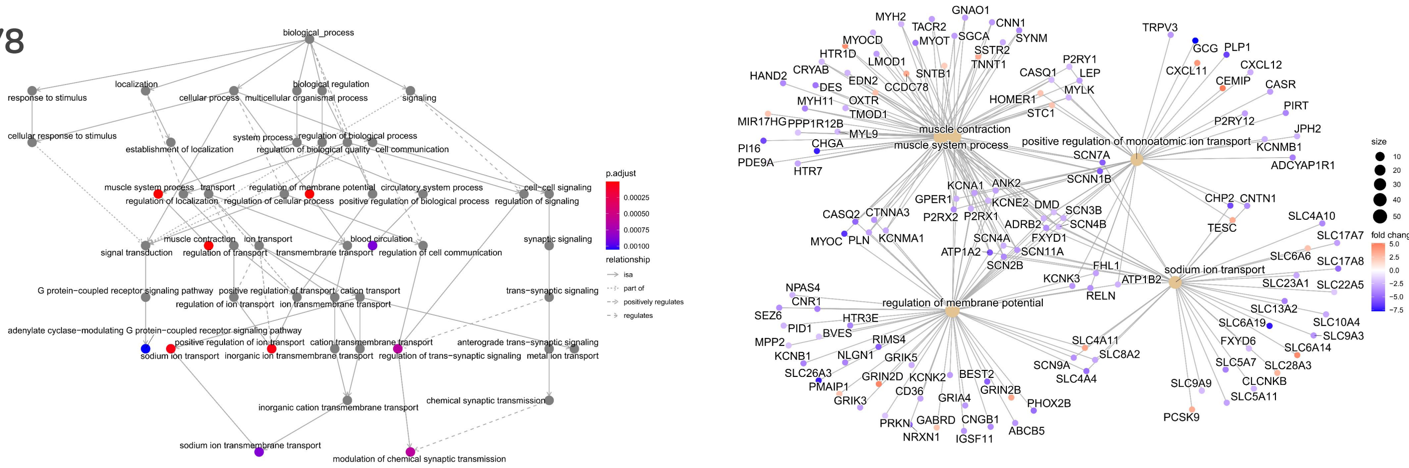

**Supplementary Figure 12. Pathway over-representation analysis for samples in DV2, DV3, Dv7 and DV8 groups.** Significant results of hypergeometric tests using as subsets differentially expressed genes between each DV group and normal group and as ontology *Biological Process* category from Gene Ontology Consortium. *Left*, significant routes (colored) displayed in a parent-child tree configuration. *Right*, differentially expressed genes members of the most significant over-represented pathways, displayed in a network configuration.
